# SELECTIVE TRANSCRIPTOMIC VULNERABILITY OF MEMBRANE-INTEGRATED ARCHITECTURES DURING NEURAL TISSUE VITRIFICATION

**DOI:** 10.64898/2026.03.26.714628

**Authors:** Dominika Wilczok, Qian Long, Zhuoan Huang, Joseph Kangas, Yewei Liu, Minyan Wang, Ferdinand Kappes

## Abstract

Cryopreservation is essential for long-term storage of biological tissues. Yet, surprisingly, the precise molecular impact of cryopreservation on tissue transcriptomes remains poorly defined. This study provides the first resource of whole-genome transcriptomic changes following cryopreservation. This study used bulk RNA sequencing to examine how preservation method (snap freezing or vitrification) affects transcriptomes in mouse cerebral cortex and hippocampus. This allowed us to separate cryoprotectant-specific changes from cold induced-changes via snap freezing. In a subset of genes, tissues processed under vitrification conditions showed selective under-representation of a small but structurally coherent group of transcripts, with the hippocampus exhibiting greater vulnerability than the cortex. UniProt annotation revealed that affected transcripts were strongly enriched for proteins with membrane-associated, secretory-pathway, and multi-pass topologies, indicating that structurally complex membrane-integrated architectures are disproportionately sensitive to vitrification. Pathway-level analysis using iPANDA further showed that negative preservation scores in vitrified tissue clustered primarily within signal transduction and metabolic pathways, suggesting coordinated pathway-level disruption rather than global transcript loss. Together, these results demonstrate that vitrification conditions induce selective and structured molecular perturbations in neural tissue, defined by the under-recovery of transcripts associated with membrane and secretory pathway organization. This work highlights molecular vulnerability during vitrification and emphasizes the need for transcript-level evaluation when optimizing cryopreservation approaches for neural systems.

## INTRODUCTION

Cryopreservation is a foundational technology in biotechnology, enabling the long-term storage of biological materials and supporting a wide range of applications in basic research, medicine, and conservation, such as routine access to viable cells for culture and transplantation, long-term preservation of tissues for molecular analysis, in vitro disease models, and genetic resource banking for regenerative medicine, biodiversity conservation, and reproductive medicine ^1^. Cryopreservation relies on cooling biological specimens to cryogenic temperatures, most commonly storage in liquid nitrogen (−196 °C) or in the vapor phase above liquid nitrogen (≈−150 °C), and in some cases in ultra-low mechanical freezers (≈−80 °C). At these temperatures, molecular diffusion and biochemical reactions are greatly reduced, effectively arresting biological activity ^2^. Depending on the goal of preservation and sample type, different cooling approaches are used. When the objective is to preserve molecular composition for downstream analyses such as proteomics or metabolomics, samples are commonly snap frozen by rapid immersion in liquid nitrogen. Although snap freezing renders the sample non-viable, it preserves its molecular composition. Rapid cooling arrests metabolic and enzymatic activity and limits ice crystal growth, allowing DNA, RNA, proteins, and metabolites to be retained in a state suitable for downstream biochemical analysis. On the other hand, for applications where structural and functional preservation is required, the samples are vitrified. Vitrification refers to the solidification of a solution into an amorphous glass without ice crystallization, achieved through sufficiently high concentrations of cryoprotective agents (CPAs) combined with rapid cooling that suppresses ice nucleation and growth. CPAs are chemical compounds added to biological samples to mitigate damage during freezing. CPAs function by reducing ice nucleation and growth, lowering the freezing point of intracellular and extracellular solutions, and stabilizing cellular structures such as membranes and proteins ^2^. CPAs are typically categorized as penetrating agents (e.g., dimethyl sulfoxide (DMSO), glycerol, ethylene glycol, propylene glycol), which permeate cell membranes and reduce intracellular ice formation, and non-penetrating agents (e.g., sucrose, trehalose, hydroxyethyl starch), which primarily exert osmotic and extracellular protective effects. However, despite substantial progress at the level of cells and small tissues, scaling cryopreservation to whole organs has thus far proven technically prohibitive, with a few notable examples, due to challenges associated with CPA delivery ^3,4^, CPA toxicity ^5–8^, ice crystal formation ^9–11^, and safe, uniform rewarming scaling nonlinearly with biological size and complexity ^12,13^. Notably, a rat kidney was successfully vitrified, rewarmed, transplanted, and shown to restore close to normal physiological function in the recipient ^14^. Similar approaches, although without transplant validation, have been reported for other organs, including rat hearts ^15^, sheep ovaries ^16^, and porcine livers ^17^, with preservation of structural integrity and partial functional outcomes. Importantly, functional or structural recovery at the organ level does not necessarily imply preservation of molecular integrity. Current organ cryopreservation strategies rely on vascular perfusion with CPA to achieve uniform distribution prior to cooling. It implicitly assumes that a single cryoprotectant formulation should preserve molecular integrity across diverse tissue types exposed simultaneously during loading, vitrification, unloading and rewarming. Heterogeneous tissue composition of a whole organism, metabolic state, and permeability, however, raises the clear possibility that cryoprotectants may differentially affect molecular preservation across organs and regions, even under identical perfusion conditions. Transcriptome-wide RNA sequencing (RNA-seq) provides a means to systematically interrogate these molecular consequences. RNA-seq enables highly sensitive detection of method-specific transcript representation or loss that may not be apparent from structural or functional assessments alone. Despite widespread use of RNA-seq in biomedical research, its application in deciphering cryopreservation-induced molecular bias remains limited.

Although cryopreservation-induced molecular damage such as protein misfolding or DNA double strand breaks may occur in any tissue, their implications are especially consequential in the brain, where preservation of molecular integrity is essential for maintaining higher-order neural function. Consequently, brain cryopreservation represents a technically demanding test case for evaluating molecular preservation strategies. Notable success cases in brain cryopreservation include the functional revival of hippocampal slices after vitrification ^18^ and successful cryopreservation and rewarming of brain organoids ^19^. However, state-of-the-art approaches were set by German *et al*. (2025), showing that basic electrophysiology can be regained in hippocampi dissected from whole vitrified mouse brains ^20^. Although the whole brain showed dehydration and darkening, the hippocampal granule cells obtained from the dissected hippocampi showed regained functional electrophysiological measures like basic excitability, E/I balance (a measure of the relative strength of excitatory versus inhibitory synaptic transmission), spontaneous activity, input–output transmission, and long-term potentiation were preserved relative to untreated control slices. The whole dissected hippocampi, however, showed reduced mitochondrial respiration and impaired short-term plasticity. The work of German and colleagues (2025) suggests that although whole-brain vitrification may introduce global injury, specific regions such as the hippocampus can still retain measurable functional capacity. Yet long-term goals in cryobiology aim for preservation methods that maintain the integrity of the entire brain, not only isolated regions. Consequently, understanding the structural and physiological differences between major brain regions is essential, as these intrinsic features may reveal region-specific susceptibilities to cryoinjury. Two brain regions of interest to neuroscience given their high neuronal density and great functional relevance are the cerebral cortex and the hippocampus (a comparison is outlined in figure 1). While cortical organization is dominated by six distinct layers (I–VI) with granular, pyramidal, and multiform layers, the hippocampus contains compact laminae such as the dentate granule cell layer and the CA pyramidal cell layer ^21,22^. Both regions contain pyramidal neurons, interneurons, and dense excitatory glutamatergic projection systems, however, the cortex contains a wide variety of projection neurons (pyramidal and spiny stellate) and at least 15 interneuron subtypes, the hippocampus contains fewer neuronal classes but extremely dense and well-defined granule and pyramidal layers ^21,22^. The cortex, compared to a hippocampus, exhibits a denser and highly organized network of perineuronal nets, which are specialized extracellular matrix structures that wrap around certain neurons (mostly parvalbumin-positive interneurons) and regulate synaptic plasticity, ion buffering, and protection against oxidative stress ^23^. Astrocytes in the cortex are highly coupled, while in the hippocampus they are only coupled in the CA1 region ^24^. Both regions have a high proportion of pericytes compared to other brain regions^24^.

**Figure 1:**
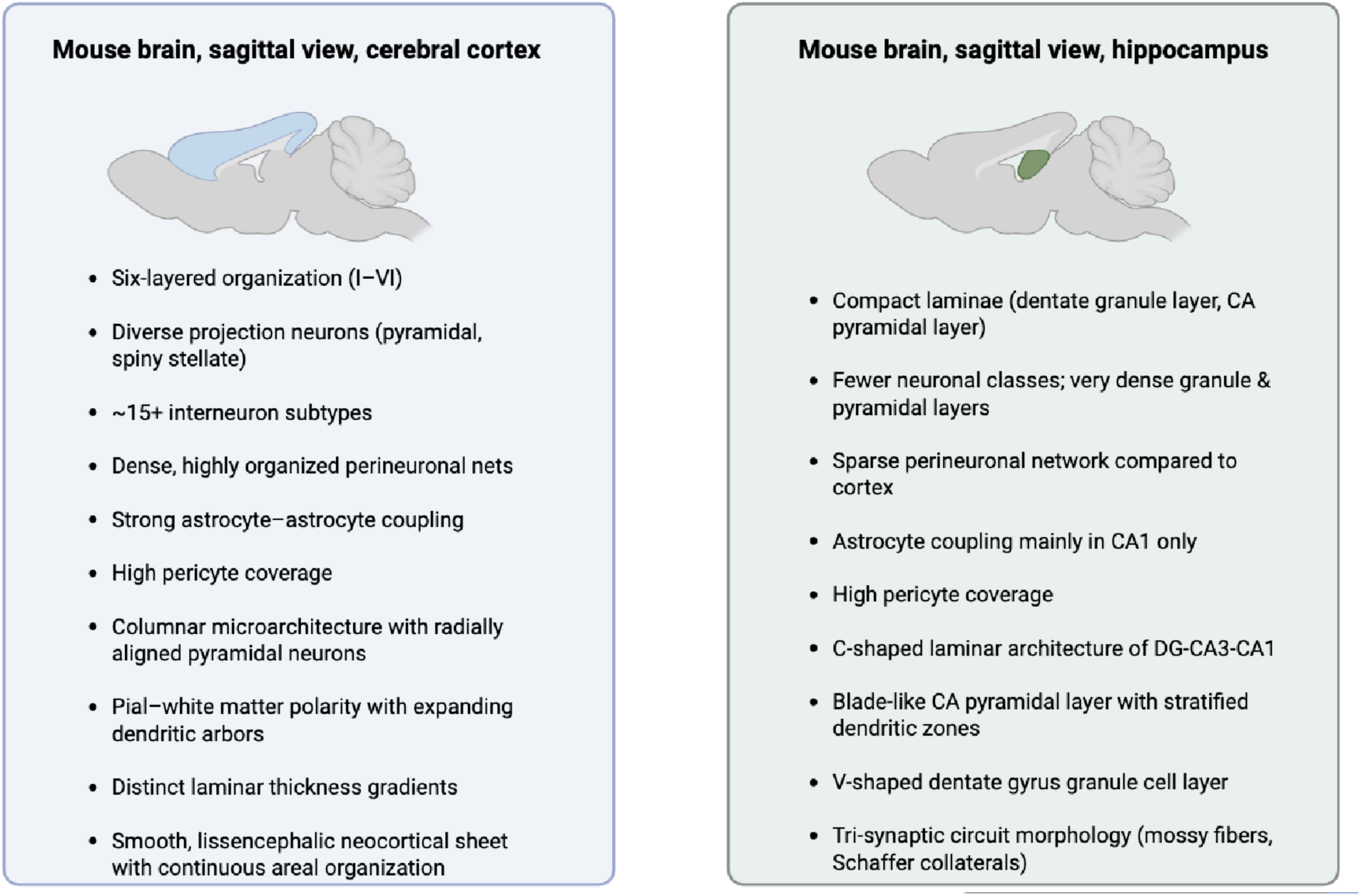
Histological comparison of murine cerebral cortex and hippocampus. Schematically shown are major differences between the murine cortex and hippocampus. See text for details.

These region-specific structural features imply that cryopreservation may not impact all neural tissues uniformly, making it necessary to examine how preservation strategies affect molecular integrity. To address these gaps, this work applies RNA sequencing to compare transcriptomic preservation following snap freezing and vitrification in neural tissue, proxied by isolated mice hippocampi and cerebral cortex. Snap freezing, which is not intended to support post-thaw viability or functional recovery but is widely used to preserve molecular content, serves here as a molecular reference condition ^25^. In contrast, vitrification has been shown to enable structural and, in select cases, functional preservation of complex tissues and organs, yet its molecular consequences remain poorly characterized ^12,17^. The analysis focuses on transcripts that are selectively under-represented after vitrification while remaining relatively stable under snap freezing, hypothesizing that such transcripts are selectively vulnerable to CPA exposure rather than cold-induced effects. The scope of the study is limited to RNA representation and does not address post-thaw viability, functional recovery, or organismal revival, but instead evaluates transcriptomic preservation with relevance for cryoprotectant and protocol optimization.

## MATERIAL AND METHODS

### Tissue acquisition

All animal procedures were performed at the Laboratory Animal Centre, Soochow University with Prof. Wang’s laboratory, under a mutual agreement between Xi’an Jiaotong-Liverpool University and Soochow University. All protocols followed the institutional animal care and use committee (IACUC) guidelines of XJTLU, with ethical approval number ER-SRR-1088616620231127132732. A trained and authorized professional euthanized three adult female mice (*Mus musculus,* C57BL/6 strain, aged 8-12 weeks) by cervical dislocation to minimize distress and ensure rapid cessation of vital functions. The hippocampus and cerebral cortex (Figure 1) were selected as tissues of interest due to their central roles in memory formation and higher-order cognitive functions, as well as their widespread use as reference regions in studies of neural integrity and molecular preservation.

Before tissue acquisition artificial cerebrospinal fluid (ACSF) was prepared from: 125 mM NaCl, 2.5 mM KCl, 1.2 mM NaH₂PO₄, 12.5 mM D-glucose, 2 mM CaCl₂, 2 mM MgSO₄, and 26 mM NaHCO₃. All salts were dissolved in ddH₂O to a final volume of 1 L, bubbled with carbogen (95% O₂ / 5% CO₂) for ∼10 min, and the pH was adjusted to ∼7.3 with <1 mL of 12 N HCl. The solution was stored at 4 °C until use.

For preparation of cerebral cortex and hippocampus samples, the following steps were performed. Each of the euthanized mice was decapitated, and the brain was carefully removed from the skull. The intact brain was immediately placed in ice-cold ACSF continuously aerated with 95% O₂/5% CO₂ and maintained on wet ice to preserve tissue integrity. The cerebral cortex and hippocampus (bilateral) were then dissected under a stereomicroscope, ensuring minimal damage to surrounding structures. Tissues were pooled to ensure that each sample contained material from all mice and were then allocated into three experimental groups: fresh, snap-frozen, or vitrified (Figure 2). Pooled samples were further split into three sequencing reactions to allow for statistical analysis.

**Figure 2:**
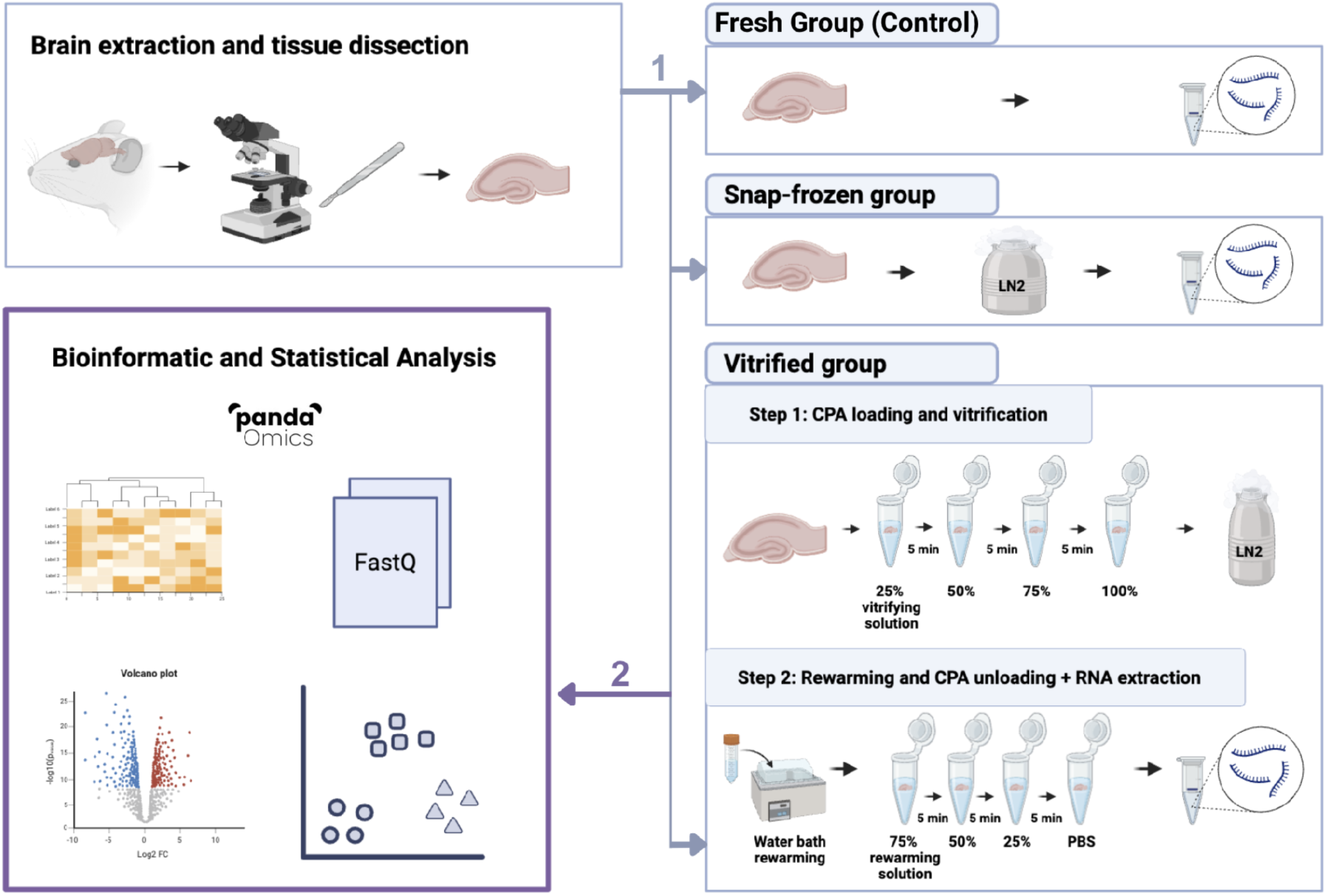
Graphical abstract of methodological procedures. The tissues extracted from the brain were equally divided into 3 groups, vitrified, snap frozen and control, as indicated. Subsequently, RNA-seq data obtained from all groups were subjected to bioinformatics data analysis.

### Control (Fresh) Group

Samples derived from the fresh condition were processed immediately for RNA extraction using Trizol (see below).

### Snap Frozen Group

For snap freezing, each sample was transferred to an Eppendorf tube and rapidly immersed in liquid nitrogen for 5 minutes. Samples were subsequently stored at −80 °C for 4 days until RNA extraction

### Vitrified Group

The vitrification stock solution (total weight of 100 wt%) was prepared by combining ethylene glycol (28 wt%) and dimethyl sulfoxide (DMSO) (22.3 wt%) as permeating cryoprotectants, sucrose (17.1 g) as a non-permeating osmotic agent, and phosphate-buffered saline (32.6 wt%) as the buffered aqueous carrier. Ethylene glycol and DMSO were used in combination to achieve a high intracellular CPA fraction required for vitrification while avoiding reliance on a single permeating agent at the full target concentration to minimize toxicity ^26^. Sucrose was included to increase extracellular osmolality and promote controlled dehydration, thereby reducing free water available for ice formation and improving stability of the vitrified amorphous phase. PBS served as the solvent and maintained buffering capacity during solution preparation and handling. The mixture was heated in a microwave for a total of 3 minutes in 1-minute intervals, with vigorous stirring between intervals to ensure complete dissolution of sucrose. Following dissolution, the stock solution was diluted with PBS to prepare working solutions of 75%, 50%, and 25% of the full strength concentration.

Hippocampal and cortical tissues were transferred to separate Eppendorf tubes and exposed to the vitrification solution in a stepwise manner to mitigate osmotic stress. As an initial pre-equilibration step, tissues were incubated in 100 μL of 25% vitrification solution for 5 minutes, after which the solution was aspirated and replaced with 100 μL of 50% vitrification solution. This exposure was intended to permit partial CPA penetration into the tissue while limiting exposure duration to reduce CPA-associated toxicity. This process was repeated with 75% and subsequently 100% vitrification solutions, with a 5-minute incubation at each step. After exposure to the 100% vitrification solution, samples containing both the tissue and the 100% vitrification solution were immersed in liquid nitrogen for 5 minutes and stored −80 °C for 4 days.

Subsequently, a rewarming stock solution (total weight of 100g) was prepared by combining 28 g ethylene glycol, 22.3 g DMSO, 34.2 g sucrose, and 15.5 g PBS. The mixture was heated in a microwave for a total of 5 minutes in 1-minute intervals, with vigorous stirring between intervals to ensure complete dissolution of sucrose. Following complete dissolution of sucrose, the stock solution was diluted with PBS to generate 75%, 50%, and 25% rewarming solutions. The sucrose concentration in the rewarming stock solution was doubled relative to the vitrifying solution to increase extracellular osmolality during CPA efflux. A higher non-permeating solute level helps counteract rehydration-driven swelling as permeating CPAs exit the tissue on warming and moderates the osmotic gradients that develop during CPA removal and reduces volume excursions that can damage multicellular structures ^27^. The stock was subsequently diluted with PBS to generate 75%, 50%, and 25% rewarming solutions for stepwise CPA unloading.

Upon removal from liquid nitrogen, the tubes with vitrified hippocampal and cortical samples were immediately rewarmed by gentle agitation in a 37 °C water bath until the vitrifying solution returned to the liquid phase. The 100% vitrification solution was then removed by pipette aspiration and replaced with 100 μL of 75% rewarming solution. Cryoprotectant unloading was performed stepwise in descending concentration order (75%, 50%, and 25%), with samples incubated for 5 minutes at each step. Following incubation in the 25% rewarming solution, tissues were submerged in 100 μL PBS for 5 minutes. RNA extraction was subsequently performed.

### Bulk RNA extraction

Following group-appropriate treatment, each sample was homogenized in TRIzol (Thermo Fisher Scientific) according to the manufacturer’s instructions using a hand-held tissue homogenizer (Qiagen). Samples were subsequently shipped to the Beijing Genomics Institute (BGI) on dry ice for RNA extraction and RNA-seq. The reference genome for transcriptomic analysis was GCF_000001635.26_GRCm38.p6. The sequencing platform was DNBSEQ and the sequencing length was PE150. The average alignment ratio of the sample comparison genome was 99.11%. The average alignment of the gene set was 75.91%; a total of 18389 genes were detected. All libraries exhibited high sequencing quality, with stable base composition and uniformly high Phred scores, supporting their suitability for downstream analyses, as shown for cortex in Supplementary Figures S1–S3 and for hippocampus in Supplementary Figures S4–S6.

### Bioinformatics Analysis

Transcriptomic data were analyzed using a combination of dedicated bioinformatics platforms and spreadsheet-based data processing. Principal component analysis and heat maps were generated using the Dr. Tom software suite provided by Beijing Genomics Institute (BGI), enabling global assessment of sample clustering and expression patterns across experimental conditions. Differential gene expression results were further examined using the PandaOmics platform developed by Insilico Medicine, which was employed to generate volcano plots and to identify and prioritize downregulated genes using the iPANDA pathway-based scoring algorithm. iPANDA computes pathway activation scores by integrating weighted expression changes across predefined pathways, applying noise-resistant normalization and penalization steps to reduce false positives and emphasize coordinated biological shifts. The resulting iPANDA-derived pathway impact metrics were used to highlight pathways disproportionately affected by preservation strategy and to rank affected gene sets accordingly ^28^.

### Statistical Analysis

All statistical comparisons of transcriptome abundance followed the thresholds defined by the sequencing provider BGI, typically using an adjusted P value < 0.05 and absolute log2 fold-change ≥ 1 unless otherwise specified. Summary statistics, data sorting, and filtering were conducted in Microsoft Excel to support downstream analyses and figure preparation. Gene annotation and functional metadata were obtained from UniProt. No imputation of missing values was performed; pathways with insufficient gene coverage were excluded from scoring.

## RESULTS

To evaluate the transcriptomic consequences of different preservation strategies, the analysis first examined global gene expression patterns across fresh, snap-frozen, and vitrified samples (Figure 2). This initial assessment aimed to determine whether preservation method and tissue type explained major sources of transcriptomic variance and to establish the overall scale and organization of preservation-induced differences before pathway-level and transcript-level investigations.

### Transcriptomic Divergence Across Preservation Methods and Tissues

For the purposes of these analyses, transcripts showing reduced abundance relative to the fresh condition are referred to as under-represented, reflecting reduced preservation or recovery rather than inferred biological downregulation. The analysis identified a substantial number of differentially expressed genes across all pairwise comparisons, with both preserved and under-represented genes observed in each condition (Figure 3a). Principal component analysis revealed clear separation between preservation conditions at the transcriptome-wide level (Figure 3b). The first principal component (PC1), explaining 96.20% of the total variance, captured the dominant differences among samples, while PC2 accounted for an additional 1.76% of the variance. Samples clustered tightly by treatment conditions, indicating high within-group consistency and a strong effect of preservation strategy on global gene expression profiles. To further examine the structure of transcriptomic differences observed across preservation conditions, hierarchical clustering heat maps were generated for differentially expressed genes in each pairwise comparison (Figure 3c-3f). These heat maps illustrate expression patterns across individual samples and provide additional resolution on the consistency of transcript representation within and between preservation methods. Heat maps are shown for vitrified versus fresh hippocampus (HV/HF) and cortex (CV/CF) (Figure 3b, d), as well as snap-frozen versus fresh hippocampus (HS/HF) and cortex (CS/CF) (Figure 3e, f). In both hippocampus and cortex, vitrification-relative comparisons showed coherent blocks of transcript under-representation and over-representation compared to fresh tissue, whereas snap-frozen samples exhibited more modest deviations. To further characterize the direction and magnitude of transcriptomic differences between preservation methods, differential expression results were visualized using volcano plots for each pairwise comparison (Figure 3g-3j). These plots integrate effect size (log₂ fold change) and statistical significance, enabling identification of transcripts exhibiting the largest changes in representation relative to the fresh condition. Comparisons involving vitrification showed a pronounced skew toward transcripts with negative log₂ fold change, whereas snap freezing resulted in fewer and more symmetrically distributed changes relative to fresh tissue.

**Figure 3.**
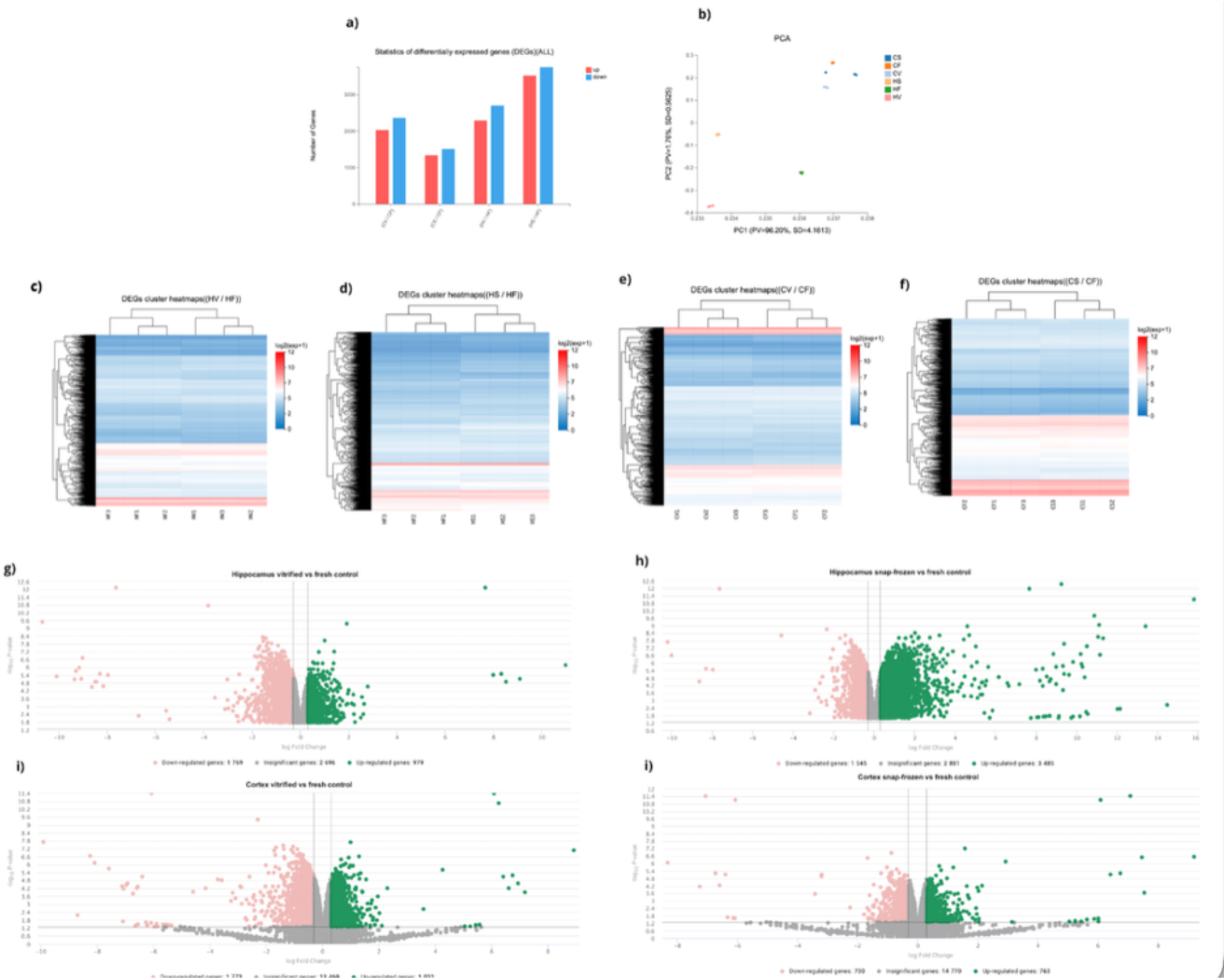
The preservation method drives global transcriptomic separation and differential gene representation. **(a)** Number of differentially expressed genes identified across all pairwise comparisons between preservation conditions. **(b)** Principal component analysis of transcriptomic profiles across all samples. C: cortex, H: hippocampus, V: vitrified, F: fresh and S: snap-frozen. **(c-f)** Hierarchical clustering heatmaps of differentially expressed genes comparing vitrified, snap-frozen, and fresh hippocampal and cortical tissues. Gene-level expression values are shown as log2(exp + 1), scaled by row, with red indicating higher expression and blue indicating lower expression relative to the gene’s mean. Dendrograms represent hierarchical clustering of both samples and genes (d) hippocampal vitrified vs hippocampal fresh. (d) hippocampal snap-frozen vs hippocampal fresh. (e) cortical vitrified vs cortical fresh. (f) cortical snap-frozen vs cortical fresh. **(g-j)** Differential transcript representation across preservation comparisons visualized by volcano plots. (g) Hippocampal vitrified vs hippocampal fresh. (h) Hippocampal snap-frozen vs hippocampal fresh. (i) Cortical vitrified vs cortical fresh. (j) Cortical snap-frozen vs cortical fresh.

### System-Level and Pathway-Level Consequences of Preservation

To determine whether transcript-level changes observed across preservation methods may cluster into coordinated biological pathways, differential expression profiles were further analyzed using the PandaOmics platform, a software suite used in biomedical research. PandaOmics’s iPANDA score integrates gene-level fold changes with pathway topology and regulatory directionality to infer pathway-level activation states, enabling assessment of higher-order structure within the transcriptomic response beyond individual genes ^28^. Subsequent analyses focused on pathways with negative iPANDA scores, with p values <0.05 as these represent pathways predicted to be inhibited relative to fresh tissue and, in the context of this study, are most consistent with reduced transcript representation associated with preservation-related effects. Each pathway with a negative iPANDA score across both tissues and conditions was filtered. In the hippocampus, iPANDA analysis identified 302 pathways with negative scores in the snap-frozen condition and 873 pathways with negative scores following vitrification (Figure 4). Of these, 210 pathways were shared between the two preservation methods. In the cortex, there were 130 pathways with a negative iPANDA score in the snap frozen cortex, and 896 negative iPANDA pathways in the vitrified hippocampus, of which 105 of these pathways overlapped.

**Figure 4:**
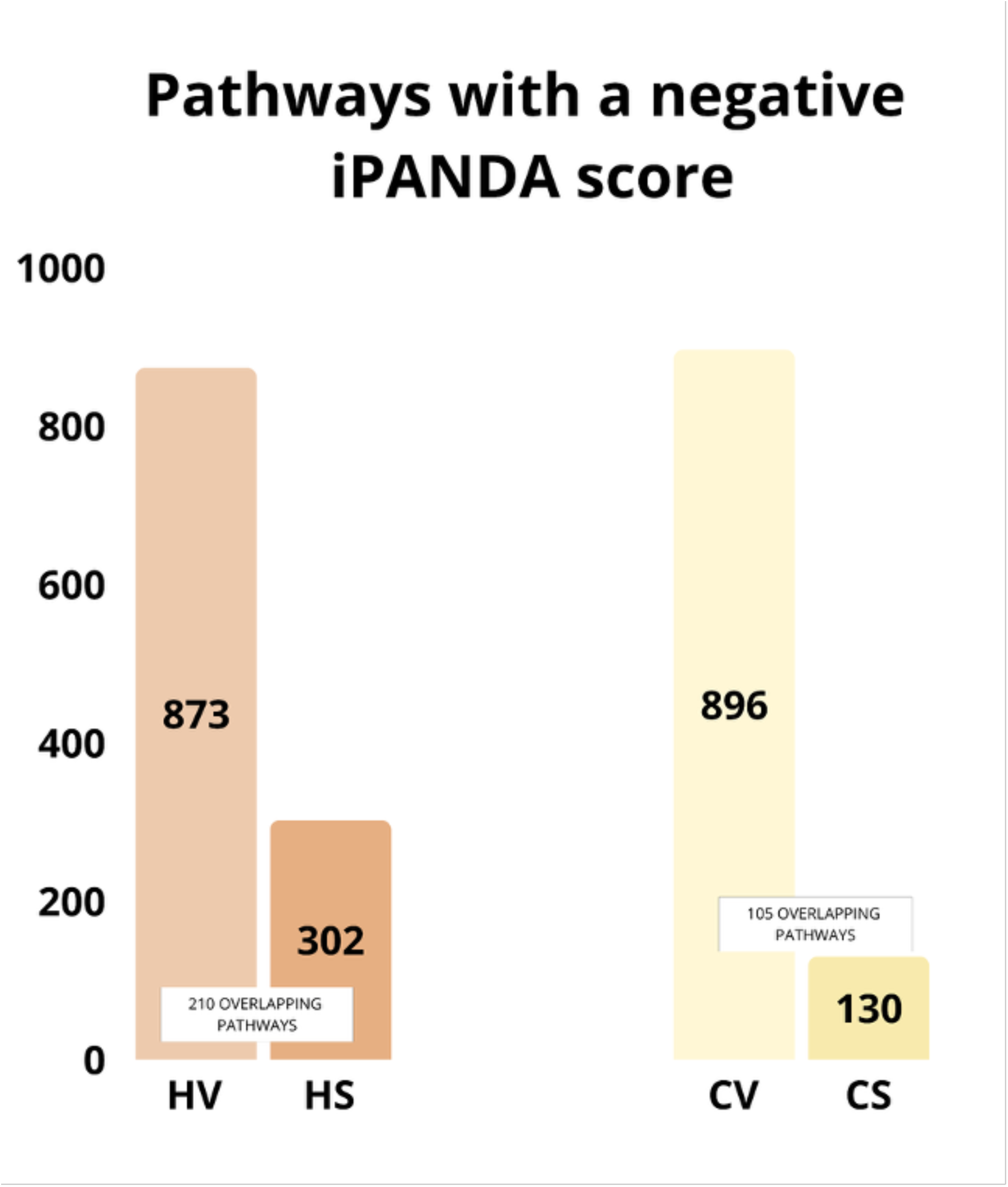
Number of pathways with a negative iPANDA scores across conditions. In the hippocampus the vitrified condition yielded nearly three-fold as many transcripts with a negative iPANDA score as the snap frozen condition (with 210 overlapping pathways between both groups). In the cortex, the vitrified condition yielded nearly seven-fold more transcripts with a negative iPANDA score than the snap frozen condition (with 105 overlapping pathways between both groups).

These differences in the magnitude and overlap of negatively scored pathways prompted a deeper assessment of how these pathways cluster into functional categories within each tissue and preservation condition. Figure 5a summarizes the number of pathways within each functional category that were shared between snap-frozen and vitrified conditions within individual tissues. Overlap was evaluated separately for hippocampus and cortex, and pathways were not required to be common across tissues. The largest numbers of shared pathways were observed in the signal transduction, metabolism, disease-related, developmental biology, and immune system categories. When this criterion was further restricted to retain only pathways shared across both preservation conditions and both tissues, the number of overlapping pathways was reduced to 13 (Figure 5b).

**Figure 5:**
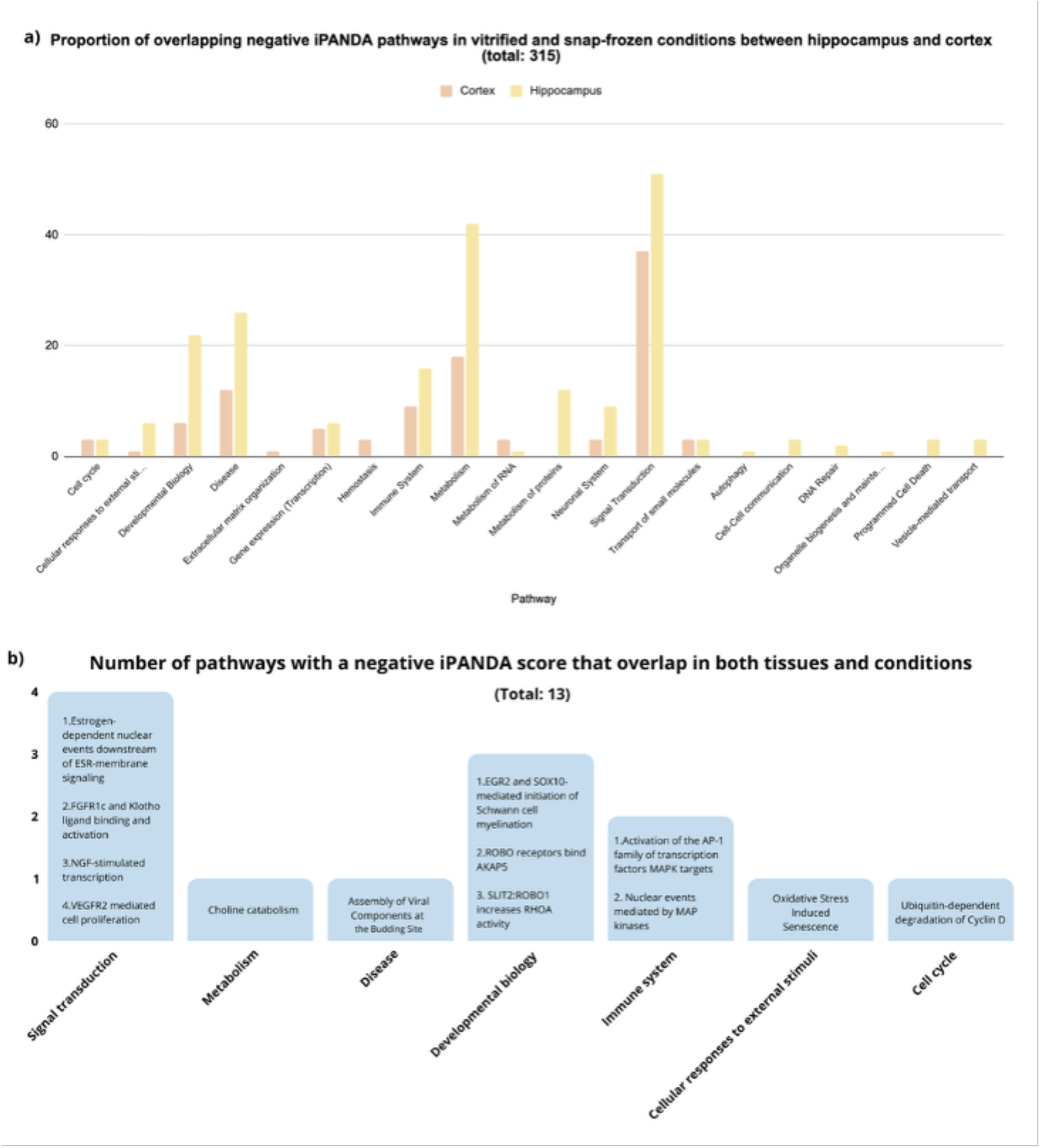
Impact of preservation method on cellular pathways. **(a)** Within-tissue pathway overlap between vitrified and snap-frozen samples **(b)** Pathways shared across tissues and preservation conditions.

To assess whether negative iPANDA pathway scores reflected coherent gene-level behavior or mixed transcript responses within pathways overlapping in both tissues and conditions, genes contributing negatively to scoring pathways were further classified based on the consistency of their directionality across preservation comparisons. Transcripts exhibiting uniform directionality across all tissues and preservation conditions either consistently positive or consistently negative log₂ fold changes were classified as consistently preserved or consistently under-represented, respectively. Transcripts showing mixed directionality, defined as a negative log₂ fold change in at least two comparisons irrespective of tissue or preservation method, were classified as bidirectionally preserved. To reduce noise from minor fluctuations, only transcripts with absolute log₂ fold-change values greater than 0.1 were included in this analysis. Transcripts exhibiting consistent directionality in only three out of four tissue–condition comparisons were excluded from this analysis to ensure stringent classification and to avoid ambiguity arising from partial or condition-specific effects. None of the transcripts contributing to any of the pathways in the categories of gene expression (transcription), metabolism of RNA, transport of small molecules or neuronal system categories met the selection criteria. Gene-level classification within negatively scoring pathways indicated that only a small fraction of transcripts exhibited consistent directionality across tissues and preservation methods, while the majority did not conform to a uniform preservation or loss pattern (Figure 6).

**Figure 6:**
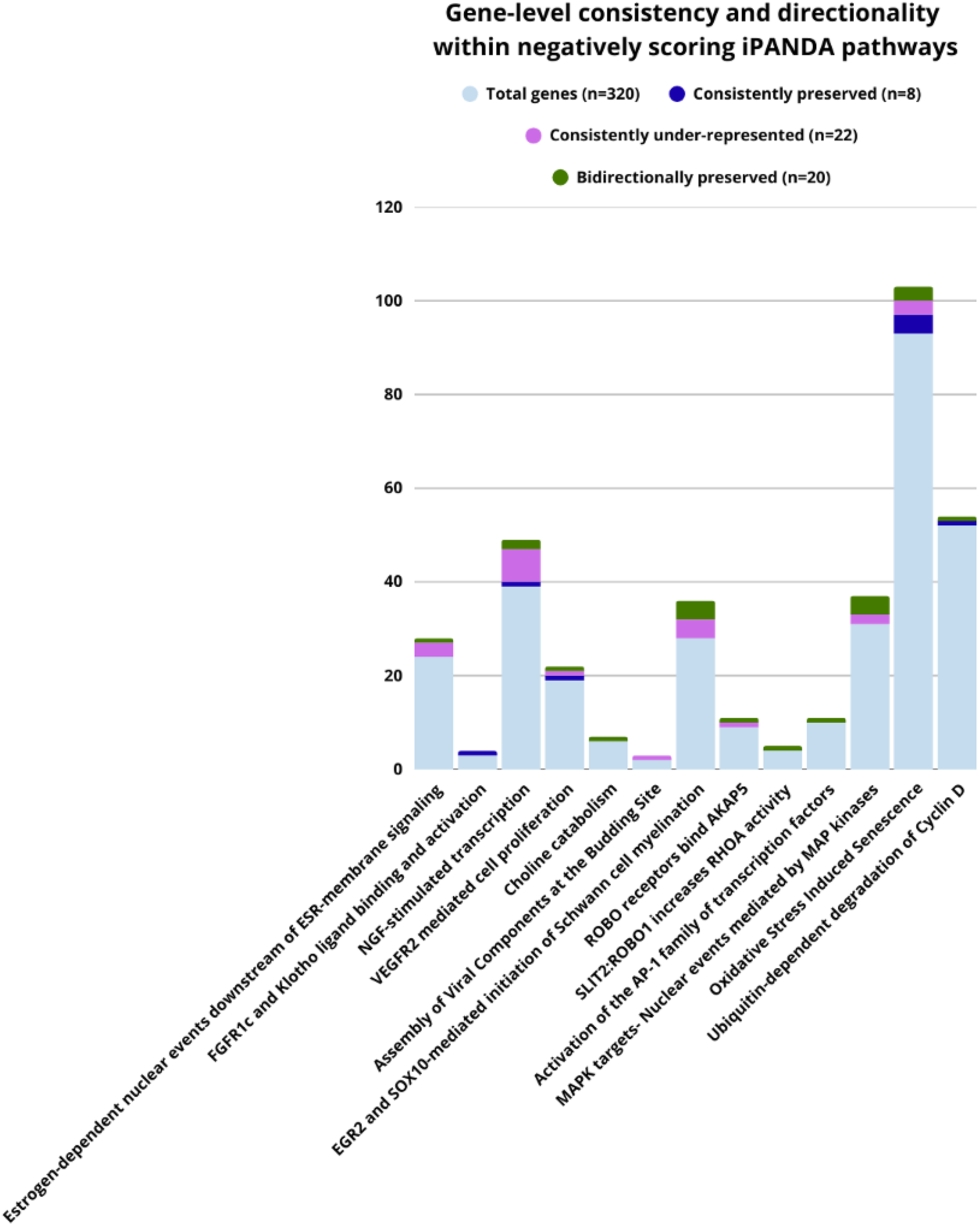
Distribution of total, consistently preserved, under-represented, and bidirectionally preserved genes within negatively scoring iPANDA pathways. Gene distribution within negatively scoring iPANDA pathways by preservation behavior. Bars represent total gene counts per functional category, partitioned into genes preserved across both methods (dark blue), consistently under-represented genes (purple), and genes with opposite directionality between tissues or preservation conditions (green).

### Selective Transcript Vulnerability to CPA exposure

Rather than pursuing individual gene examples, the analysis further focused on transcript abundance patterns and preservation logic. Transcripts that remain comparable between fresh and snap-frozen tissue but are markedly under-represented following vitrification were hypothesized to represent transcripts selectively vulnerable to cryoprotectant exposure rather than cold-induced effects. To operationalize this preservation logic at the transcript level, log₂ fold-change values were computed for all detected transcripts for each tissue and preservation method, including log₂(CS/CF) and log₂(CV/CF) for cortex, and log₂(HS/HF) and log₂(HV/HF) for hippocampus. Transcripts were then filtered independently within each tissue to identify those showing minimal deviation between snap-frozen and fresh samples, alongside pronounced under-representation following vitrification. Specifically, transcripts were required to exhibit near-zero-fold change under snap freezing relative to fresh tissue and a substantial negative fold change under vitrification, consistent with selective vulnerability to cryoprotectant exposure rather than cold-induced effects. Following tissue-specific filtering, overlap analysis was performed to identify transcripts meeting these criteria in both hippocampus and cortex (Figure 7a).

**Figure 7:**
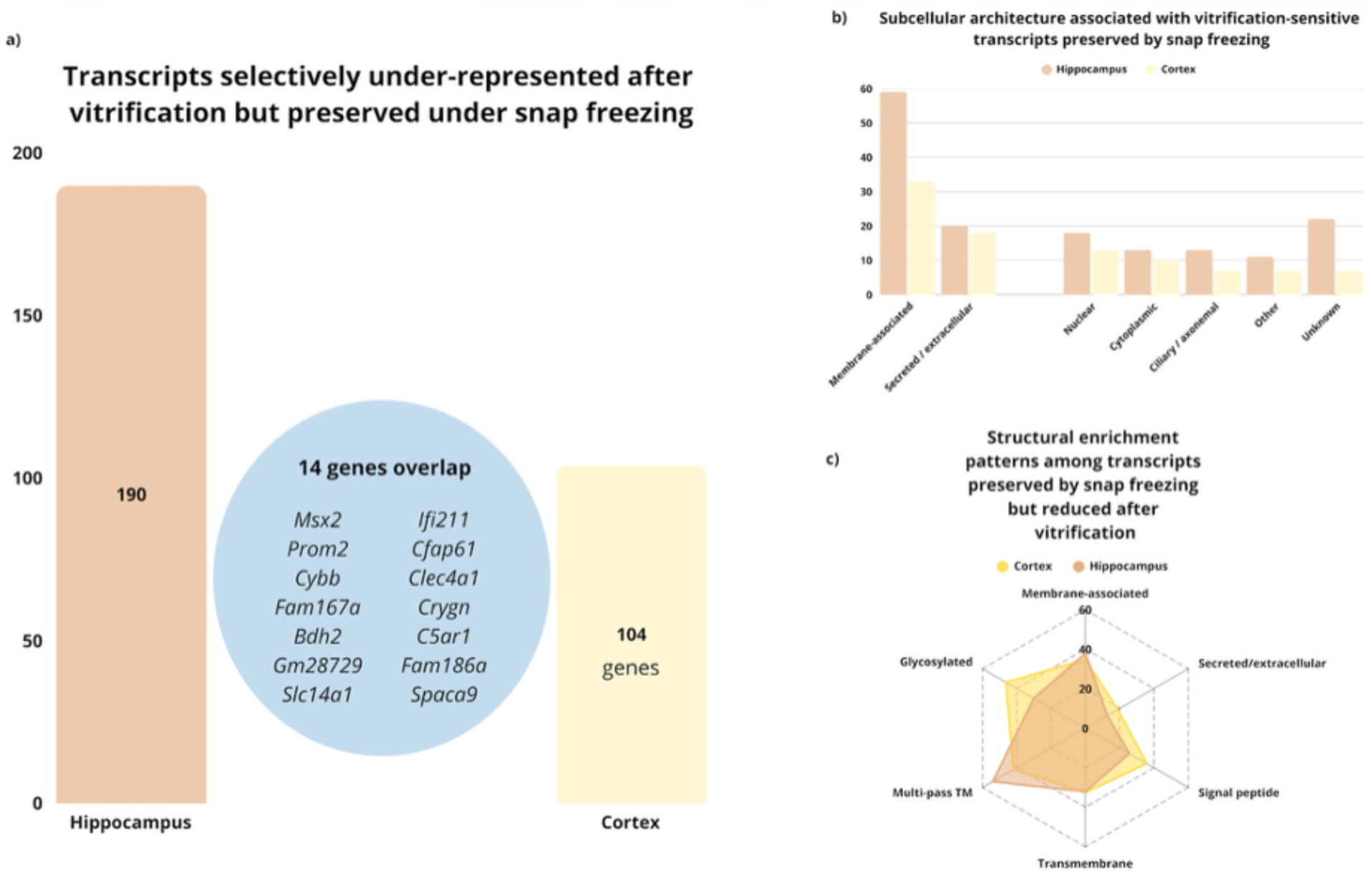
Functional characterization of altered transcripts. **a)** Transcripts selectively under-represented after vitrification but preserved after snap freezing. In the hippocampus, 190 transcripts were uniquely under-represented, whereas 104 transcripts were uniquely under-represented in the cortex. The central overlap indicates the 14 genes that exhibited vitrification-specific under-representation in both regions, suggesting shared susceptibility across tissues. **b)** Preservation-dependent changes across brain regions and subcellular architecture. **c)** Subcellular architecture and structural enrichment in snap-frozen versus vitrified samples.

To assess whether transcripts that were selectively reduced after vitrification shared common structural features, we examined the annotated subcellular localization of the proteins they encode using UniProt annotations in both tissues (Figure 7b). UniProt successfully mapped 95 of 104 cortex genes and 156 of 190 hippocampus genes. In both regions, vitrification-sensitive transcripts showed a strong and consistent enrichment for proteins routed through the secretory pathway. In the cortex, 33 of 95 genes (34.7%) encoded membrane-associated proteins and 18 (18.9%) encoded secreted or extracellular proteins, such that 53.6% of the set corresponded to surface-exposed or extracellularly trafficked protein architectures. Hippocampus-derived samples showed a very similar pattern, with 59 of 156 genes (37.8%) encoding membrane-associated proteins and 20 (12.8%) encoding secreted proteins, yielding a combined 50.6% representation of secretory-pathway targets. In contrast, classical intracellular proteins—including nuclear and cytosolic species—accounted for only 24.2% of cortex transcripts (23/95) and 19.8% of hippocampal transcripts (31/156). Transcripts encoding proteins localized to specialized structural compartments such as cilia and axonemes were also consistently overrepresented (7.4% in cortex and 8.3% in hippocampus), indicating a bias towards non-cytosolic, architecturally complex compartments. In contrast, protein (transcript) length distributions did not reveal a dominant size class and showed no evidence that extreme protein lengths drive transcript loss. In the cortex, proteins clustered primarily within a mid-length range: 62 of 95 proteins (65.3%) were 200–600 amino acids long. Hippocampus displayed the same pattern, with 95 of 156 proteins (60.9%) falling in the mid length category. Beyond subcellular localization and protein size, additional structural annotations revealed further convergence on secretory-pathway and membrane-associated architectures (Figure 7c). Signal peptides were present in 35.8% of cortex proteins (34/95) and 25.6% of hippocampal proteins (40/156), and transmembrane segments were identified in 32.6% (31/95) and 32.1% (50/156), respectively. When considered together, more than half of the transcripts in each tissue encoded proteins requiring either signal-peptide-directed ER entry, membrane insertion, or both (54.7% in cortex; 42.9% in hippocampus). Hippocampus displayed a disproportionate enrichment of multi-pass membrane proteins, with 27 of 50 transmembrane proteins (54.0%) containing two or more membrane-spanning helices, including a prominent cluster of 7-TM architectures characteristic of GPCR-like or transporter-like topologies. Cortex had 31 total transmembrane proteins, of which 13/31 were multipass. Glycosylation patterns reinforced this theme: 47 of 156 hippocampal proteins (30.1%) carried annotated glycosylation sites—primarily N-linked—and most of these were simultaneously membrane-associated or signal-peptide–containing (70.2% and 59.6%, respectively), consistent with proteins that depend on ER/Golgi processing falling between 200 and 600 amino acids.

## DISCUSSION

Cryopreservation imposes multiple biophysical and biochemical stresses on biological systems, yet the transcriptome-wide consequences of different preservation strategies remain incompletely understood, particularly in structurally complex neural tissues. This study identified a selective set of transcripts that were preserved under snap freezing but under-represented following vitrification in both murine cortex and hippocampus. Although this group represents only a small fraction of the ∼17000 detected transcripts, its composition was highly non-random. Affected genes were consistently enriched for membrane-associated proteins, multi-pass transporters, secreted factors, and signaling components. The limited scale of differential expression observed here is consistent with prior transcriptomic studies of cryopreserved multicellular systems. Anjos et al. found that only 2–3% of *Crassostrea angulata* larval transcripts changed significantly after freeze–thawing despite clear physiological impairment, with effects concentrated in membrane biology, receptor-mediated signaling, and intracellular trafficking rather than diffuse transcriptomic disruption^29^. Similarly, Cordeiro *et al*. demonstrated that vitrification of bovine oocytes produced structured, pathway-specific transcriptomic shifts enriched for glycoproteins, transporters, and membrane-associated signaling molecules even when cryoprotectant exposure alone produced minimal alterations ^30^. Together with our data, these findings point towards specific vulnerability affecting membrane-integrated and extracellular signaling architectures across diverse taxa, tissues, and cryopreservation strategies. A mechanistic basis for this recurring pattern is biologically plausible. Proteins requiring the ER-Golgi secretory system like multi-pass channels, GPCR-like receptors, adhesion molecules, and secreted factors depend on complex co-translational folding, glycosylation, lipid interactions, and stable membrane microdomains. CPAs, particularly during loading and rewarming, alter membrane fluidity, perturb lipid phase behavior, and induce osmotic compression and expansion cycles that mechanically stress these compartments ^31,32^. These volume changes mechanically stress intracellular membrane systems and may lead to membrane rupture, self-adhesion, and other structural distortions. In parallel, prolonged exposure to highly concentrated CPA solutions can produce direct toxic effects, which are often greatest at these peak-concentration stages; during unloading, the rise in temperature may further accelerate these injurious chemical and biophysical processes. Thus, the membrane-biased gene loss observed here likely reflects intrinsic biophysical sensitivities of secretory and membrane-bound architectures rather than tissue-specific artifacts. Although comparable transcriptome-wide vitrification studies in mammalian brain tissue are lacking, there are studies demonstrating feasibility of reversible cryopreservation in microglia^33^, small-molecule neural precursor cells^34^, and brain organoids ^19^. Based on previous cryoprotectant toxicity resistance research^35^, genes identified as conferring to resistance were compared with this dataset (e.g., *Myh9, Nrg2, Pura, Fgd2, Pim1, Opa1, Hes1, Hsbp1l1, Ywhag*). No coordinated pattern emerged, and effects were generally small and inconsistent across contrasts. This suggests that the selective under-representation reported here reflects a preservation-outcome axis distinct from pathways identified through forward genetic screens in cell culture. Finally, in this dataset a subset of transcripts appeared increased after vitrification relative to snap-frozen and fresh tissue. Without direct measures of transcriptional activity, we interpret these changes cautiously. They may reflect differential RNA stability in high CPA concentrations, selective depletion of other transcripts, or method-specific biases in extraction or library preparation rather than *bona fide* induction. *De novo* transcription during CPA exposure or rewarming cannot be excluded but remains speculative.

Although comprehensive transcriptomic profiling of mammalian neural tissue following vitrification remains limited, ischemic injury serves as a valuable pathophysiological proxy for understanding molecular vulnerability due to shared mechanisms of metabolic crisis and membrane disruption. In both settings, the hippocampus exhibits heightened sensitivity compared to the relatively resilient cortex; in ischemic models, this manifests as acute excitotoxic death in CA1 pyramidal neurons and abundant transcriptional dysregulation, whereas cortical populations sustain significantly milder atrophy and fewer gene expression changes ^36,37^. This established biological vulnerability parallels the molecular susceptibility observed in vitrified tissue, where the hippocampus displays a more profound under-representation of transcripts than the cortex. Furthermore, the specific functional classes compromised by vitrification, predominantly membrane-associated architectures, signal transduction components, and metabolic networks, mirror the primary axes of dysregulation in ischemia, which is characterized by the breakdown of ion channel homeostasis, the disruption of calcium signaling pathways, and maladaptive metabolic shifts such as upregulated ribosome biogenesis ^36,38^. Thus, while ischemia represents a biological reaction to injury and vitrification represents a biophysical stress, both conditions reveal that complex membrane-integrated systems and high-demand metabolic pathways constitute the pathogenic centers of neuronal vulnerability

### Limitations

Several limitations of this study should be acknowledged. First, this analysis is restricted to bulk RNA sequencing and does not distinguish between cell-type-specific effects or regional heterogeneity within tissues. Second, transcript representation was assessed after a fixed storage duration and a single vitrification protocol, limiting generalizability across cryoprotectant formulations, loading rates, or rewarming strategies. Given that transcriptional responses are among the earliest and most dynamic molecular changes triggered by cellular stress, RNA profiles offer a sensitive first layer for detecting preservation-related perturbations. Accordingly, this study focused on transcriptomics as an initial readout, with proteomic, epigenomic, and functional assessments reserved for future work. Finally, the use of pathway inference tools, while valuable for identifying higher-order structure, introduces dependence on curated pathway definitions that may not fully capture preservation-induced molecular artifacts.

### Future directions

Future studies should determine whether the transcriptomic signatures identified here correspond to functional preservation outcomes. This will require coupling RNA-seq with viability and physiological assays in vitrified neural tissue, including electrophysiology, mitochondrial function, and synaptic activity. Achieving functionality in hippocampal slices after vitrification has been previously demonstrated ^20^. Systematic evaluation of individual cryoprotectants and defined cocktails will be essential to establish whether specific CPA chemistries can mitigate the pronounced membrane- and secretory-pathway vulnerability observed in this study. Proteomic profiling will further clarify whether transcript under-representation translates into proportional protein loss or whether post-transcriptional buffering preserves functional architecture. Finally, if vitrified neural tissue can be rendered viable, longitudinal analyses after rewarming will be critical to distinguish transient CPA-loading responses from persistent molecular injury. Together, these directions will define the mechanistic basis of vitrification-induced transcript loss and guide the design of preservation strategies with improved molecular fidelity.

Despite the limitations, the findings have important implications for cryopreservation research, particularly in the context of organ-scale and whole-organism strategies that rely on vascular perfusion with a single cryoprotectant formulation. The assumption that a given vitrification protocol preserves transcriptomal integrity uniformly across tissues is not supported by the transcriptomic patterns observed here. Instead, the data suggest that cryoprotectant exposure can introduce selective molecular loss that varies across transcripts, tissues, and pathways, even when gross structural preservation is achieved. For applications where downstream molecular analysis, omics-based profiling, or long-term informational preservation is a priority, these effects warrant explicit consideration.

## CONCLUSIONS

Organ cryopreservation research is advancing rapidly, yet its effects on transcriptomic integrity remain largely unexplored. Our data show that vitrification conditions are associated with selective under-representation of transcripts encoding membrane-integrated and secretory-pathway proteins, revealing a dimension of molecular vulnerability that is not detectable through morphology or functional assays alone. Integrating molecular readouts into cryoprotectant development and protocol optimization may therefore be essential for advancing cryopreservation toward applications that demand not only survival or function but faithful preservation of biological information.

## Funding

This project is supported by DKU Office of Academic Services through a Signature Work Research Grant and additional funding by the Signature Fund.

## Declaration of Interests

The authors do not have any declarations of interest to disclose.

## Supplementary Information

Document S1

figures S1-S6

## Notes

https://www.ncbi.nlm.nih.gov/geo/query/acc.cgi?acc=GSE325022

### Competing Interest Statement

The authors have declared no competing interest.

