## Supplemental Figures S1-S6 for "SELECTIVE TRANSCRIPTOMIC VULNERABILITY OF MEMBRANE-INTEGRATED ARCHITECTURES DURING NEURAL TISSUE VITRIFICATION"

### Supplementary Information

#### RNA-seq Quality Control Metrics for All 18 Samples

Comprehensive quality control metrics for all eighteen RNA-seq libraries are provided in this appendix. For each sample, the following metrics are reported: the distribution of raw, adapter-contaminated, and low-quality reads, together with base-composition profiles and Phred quality scores of clean reads. Across all conditions and tissues, the libraries displayed stable nucleotide proportions and uniformly high base quality along the read length, with negligible adapter or low-quality contamination. These results indicate that all sequencing datasets meet the requirements for robust downstream alignment and quantitative transcriptome analysis.

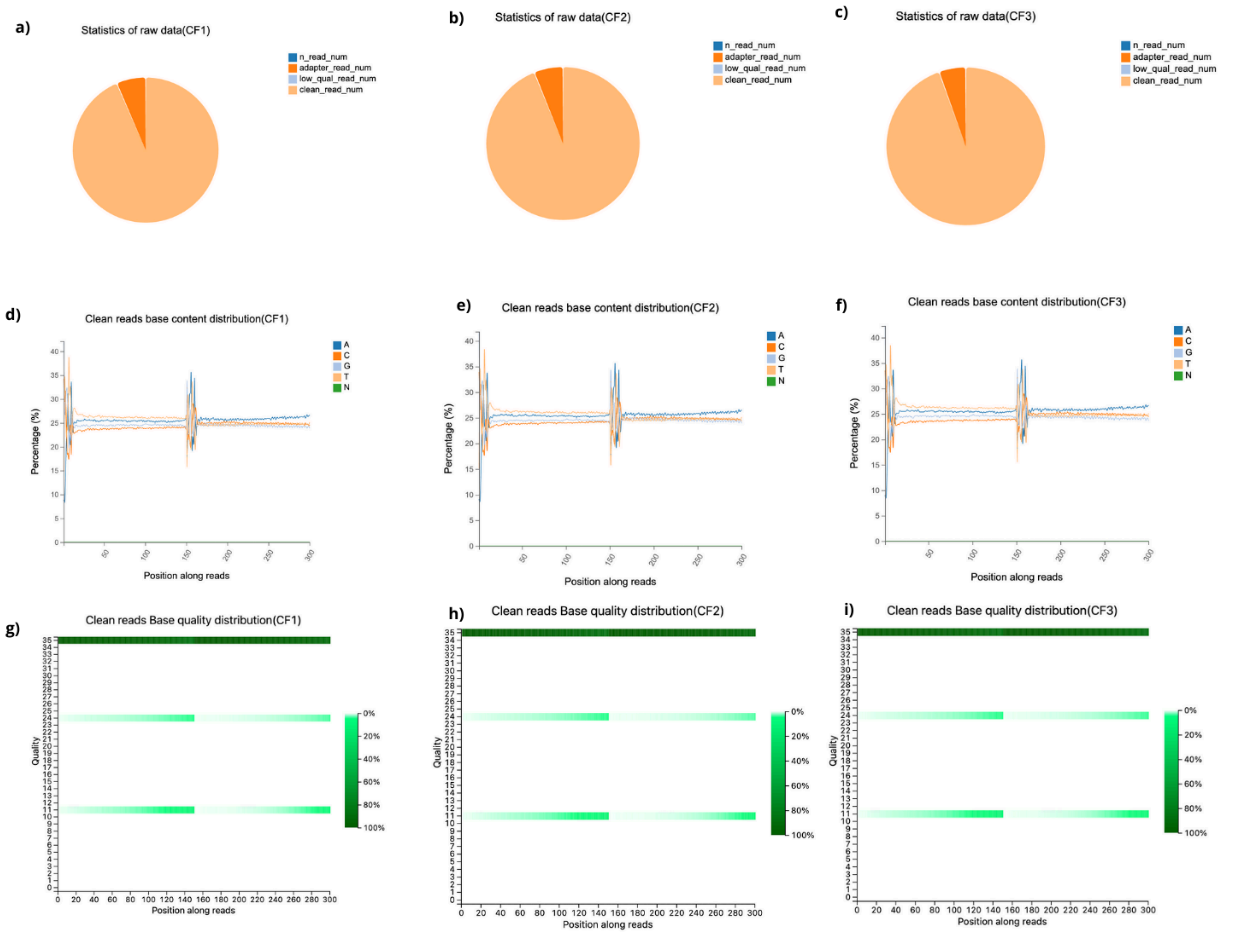


**Figure S1: Quality control metrics for fresh (control) cerebral cortex RNA-seq samples.**

(a–c) Statistics of raw sequencing data for samples CF1–CF3, showing total reads, adapter-contaminated reads, low-quality reads, and the number of clean reads retained after filtering.

(d–f) Base-content distribution along clean reads for CF1–CF3. The profiles show balanced A/T and C/G proportions across read positions without positional bias, indicating uniform nucleotide composition after trimming.

(g–i) Clean-read base-quality distributions (Phred scores) for CF1–CF3. High, stable quality scores are maintained across read length, confirming that all samples meet requirements for downstream alignment and expression quantification.

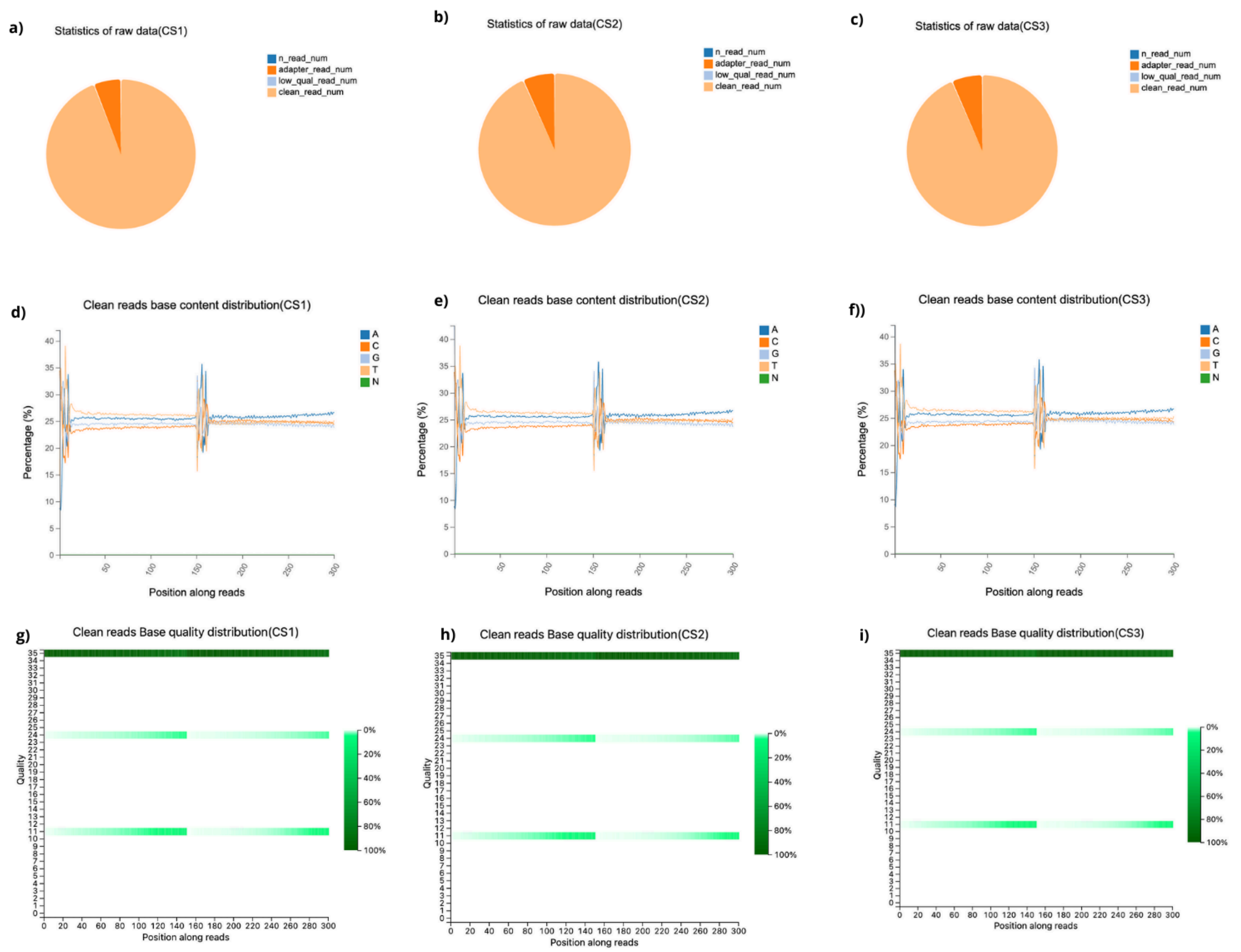


**Figure S2: Quality control metrics for snap frozen cerebral cortex RNA-seq samples.**

(a–c) Statistics of raw sequencing data for samples CS1–CS3, showing total reads, adapter-contaminated reads, low-quality reads, and the number of clean reads retained after filtering.

(d–f) Base-content distribution along clean reads for CS1–CS3. The profiles show balanced A/T and C/G proportions across read positions without positional bias, indicating uniform nucleotide composition after trimming.

(g–i) Clean-read base-quality distributions (Phred scores) for CS1–CS3. High, stable quality scores are maintained across read length, confirming that all samples meet requirements for downstream alignment and expression quantification.

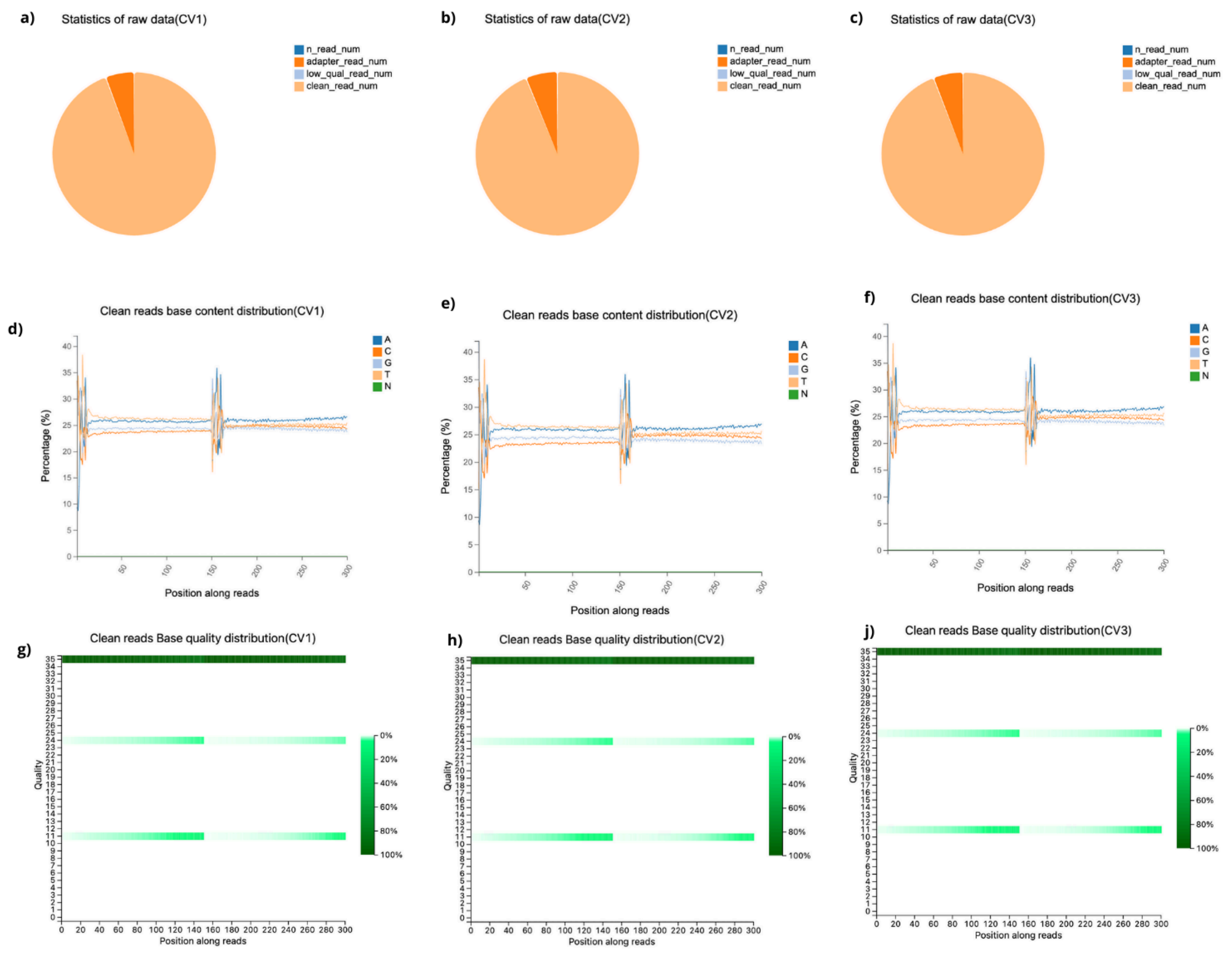


**Figure S3: Quality control metrics for vitrified cerebral cortex RNA-seq samples.**

(a–c) Statistics of raw sequencing data for samples CV1–CV3, showing total reads, adapter-contaminated reads, low-quality reads, and the number of clean reads retained after filtering.

(d–f) Base-content distribution along clean reads for CV1–CV3. The profiles show balanced A/T and C/G proportions across read positions without positional bias, indicating uniform nucleotide composition after trimming.

(g–i) Clean-read base-quality distributions (Phred scores) for CV1–CV3. High, stable quality scores are maintained across read length, confirming that all samples meet requirements for downstream alignment and expression quantification.

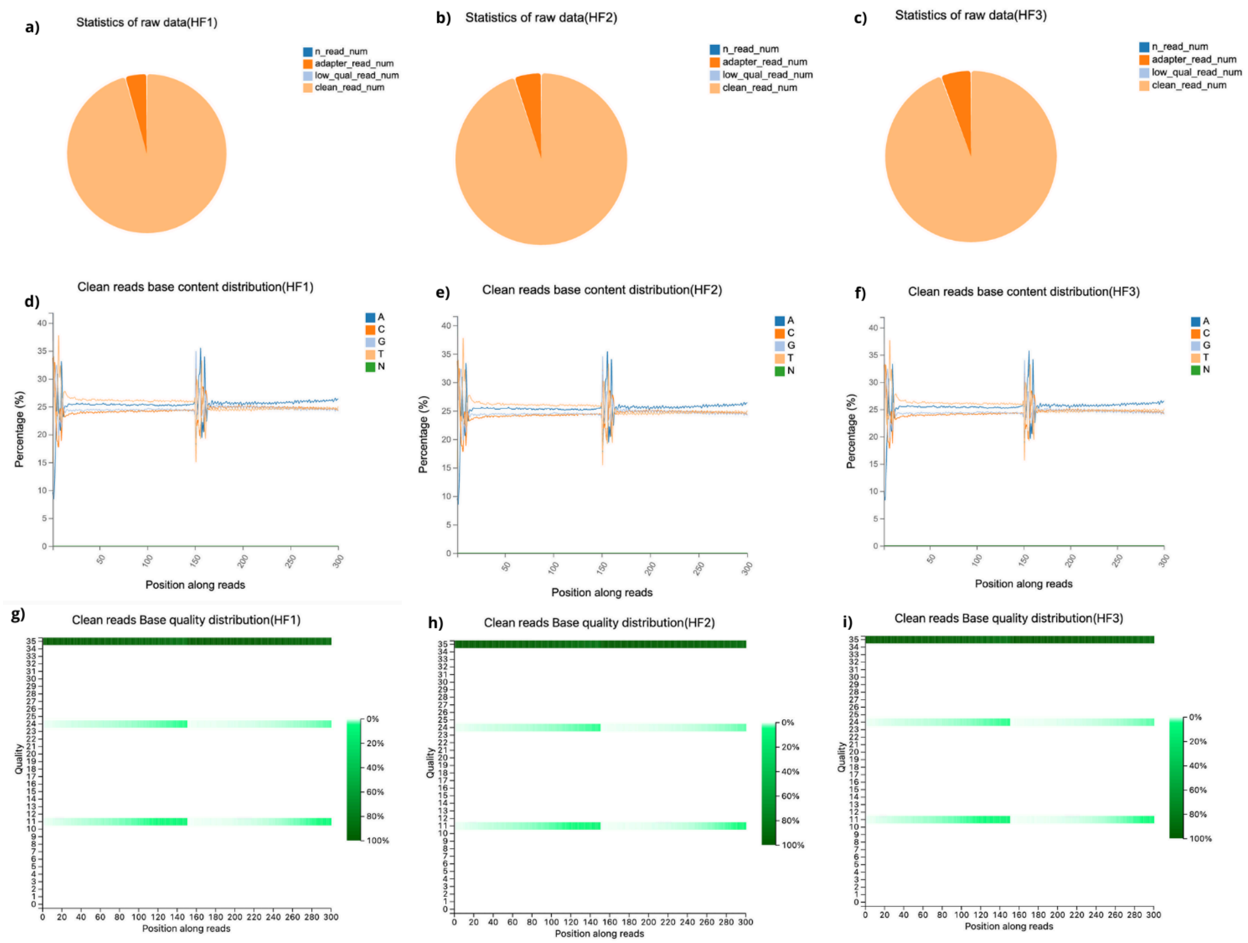


**Figure S4: Quality control metrics for fresh (control) hippocampus RNA-seq samples.**

(a–c) Statistics of raw sequencing data for samples HF1–HF3, showing total reads, adapter-contaminated reads, low-quality reads, and the number of clean reads retained after filtering.

(d–f) Base-content distribution along clean reads for HF1–HF3. The profiles show balanced A/T and C/G proportions across read positions without positional bias, indicating uniform nucleotide composition after trimming.

(g–i) Clean-read base-quality distributions (Phred scores) for HF1–HF3. High, stable quality scores are maintained across read length, confirming that all samples meet requirements for downstream alignment and expression quantification.

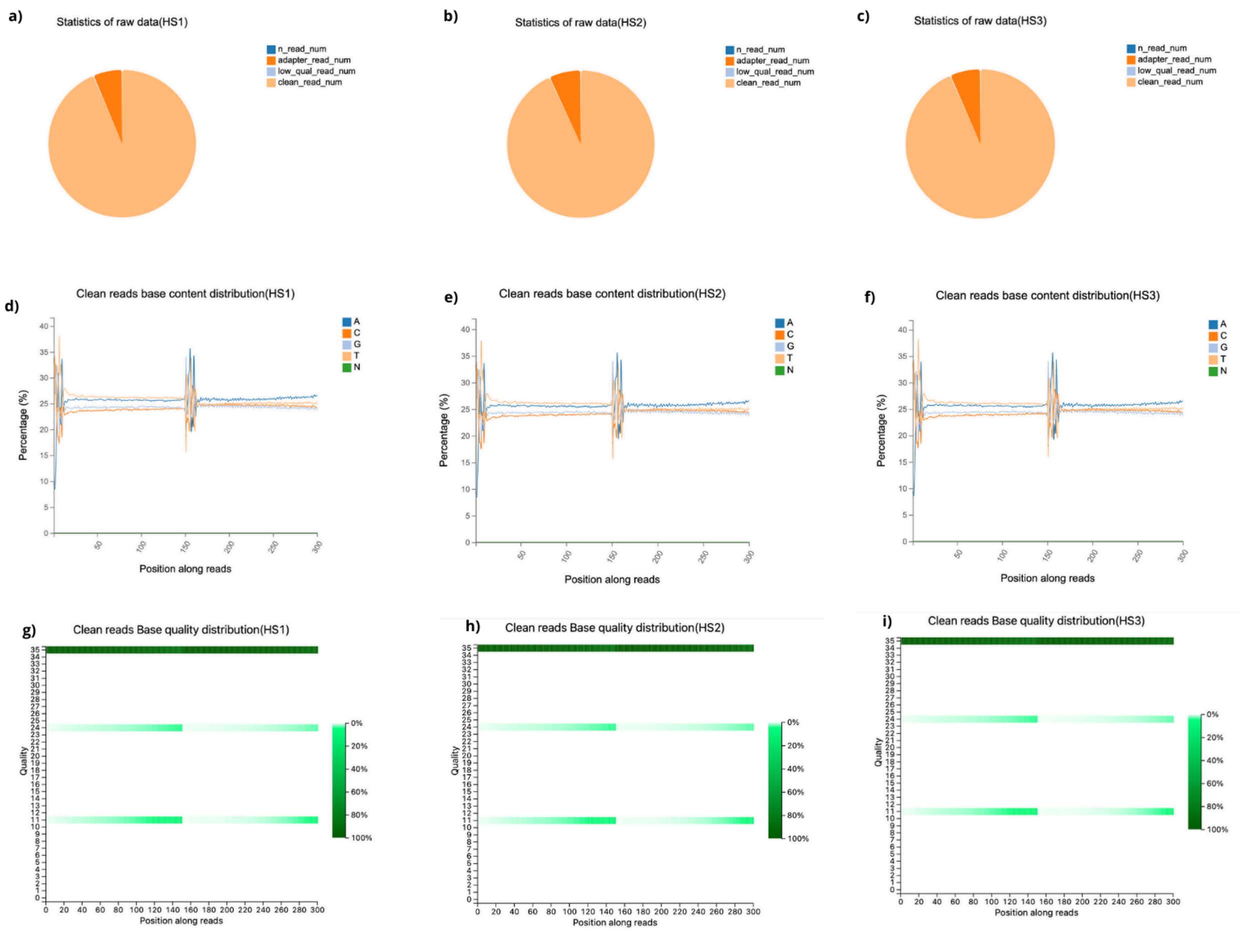


**Figure S5: Quality control metrics for snap frozen hippocampus RNA-seq samples.**

(a–c) Statistics of raw sequencing data for samples HS1–HS3, showing total reads, adapter-contaminated reads, low-quality reads, and the number of clean reads retained after filtering.

(d–f) Base-content distribution along clean reads for HS1–HS3. The profiles show balanced A/T and C/G proportions across read positions without positional bias, indicating uniform nucleotide composition after trimming.

(g–i) Clean-read base-quality distributions (Phred scores) for HS1–HS3. High, stable quality scores are maintained across read length, confirming that all samples meet requirements for downstream alignment and expression quantification.

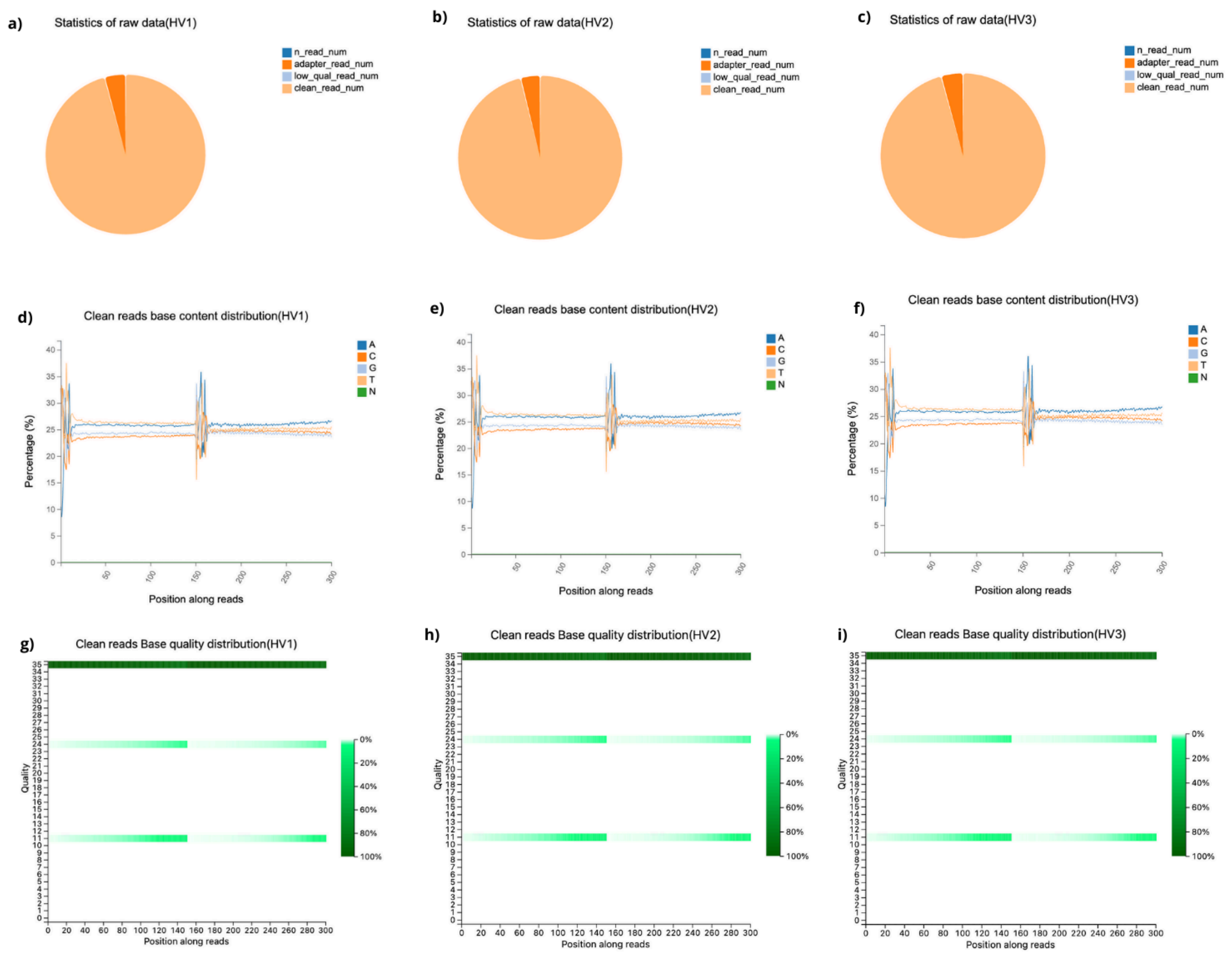


**Figure S6: Quality control metrics for vitrified hippocampus RNA-seq samples.**

(a–c) Statistics of raw sequencing data for samples HV1–HV3, showing total reads, adapter-contaminated reads, low-quality reads, and the number of clean reads retained after filtering.

(d–f) Base-content distribution along clean reads for HV1–HV3. The profiles show balanced A/T and C/G proportions across read positions without positional bias, indicating uniform nucleotide composition after trimming.

(g–i) Clean-read base-quality distributions (Phred scores) for HV1–HV3. High, stable quality scores are maintained across read length, confirming that all samples meet requirements for downstream alignment and expression quantification.
